# p75NTR upregulation following perinatal hypoxia leads to deficits in parvalbumin-expressing GABAergic cell maturation, cortical activity and cognitive abilities in adult mice

**DOI:** 10.1101/2024.01.16.574659

**Authors:** Bidisha Chattopadhyaya, Karen K.Y. Lee, Maria Isabel Carreño-Muñoz, Andrea Paris- Rubianes, Marisol Lavertu-Jolin, Martin Berryer, Frank M. Longo, Graziella Di Cristo

## Abstract

Children who experienced moderate perinatal hypoxia are at risk of developing long lasting subtle cognitive and behavioral deficits, including learning disabilities and emotional problems. Understanding the underlying mechanisms is an essential step for designing targeted therapy.

Fast-spiking, parvalbumin-positive (PV) GABAergic interneurons modulate the generation of gamma oscillations, which in turn regulate many cognitive functions including goal-directed attentional processing and cognitive flexibility. Due to their fast firing rate, PV cell function requires high levels of energy, which may render them highly vulnerable to conditions of metabolic and oxidative stress caused by perinatal hypoxia. Here, we show that adult mice that experienced moderate perinatal hypoxia (MPH) have decreased cortical PV expression levels in addition to specific impairments in the social, recognition memory and cognitive flexibility domain. We further found that the expression level of the neurotrophin receptor p75NTR, which limits PV cell maturation during the first postnatal weeks, is increased in MPH mice. Genetic deletion of p75NTR in GABAergic neurons expressing the transcription factor Nkx2.1, which include PV cells, protects mice from PV expression loss and the long-term cognitive effects of MPH. Finally, one week treatment with a p75NTR inhibitor starting after MPH completely rescues the cognitive and cortical activity deficits in adult mice. All together this data reveals a potential molecular target for the treatment of the cognitive alterations caused by MPH.

## Introduction

Despite the important advances in perinatal care in the past decades, neonatal hypoxia remains a severe condition leading to behavioral alterations and cognitive deficits (Saliba et al, 1999; American College of Obstetrics and Gynecology, 2003; Volpe, 2008). The large majority of efforts to understand how hypoxia affects brain development have been devoted to create experimental models of severe hypoxia or hypoxia-ischemia, with a focus on lesion site and size (Roberton et al, 1988, 1989). In fact, it was thought that absence of cerebral palsy or other major disabilities at a young age indicated a good long-term prognosis (Barnett et al, 2002). Much less is known regarding the effects of moderate hypoxia on long-term brain development and cognition. Recently, it has been reported that cognitive problems in patients that suffered mild to moderate hypoxia often do not become visible until school age and that these troubles can occur in absence of motor deficits (Dilenge et al, 2001; Maneru et al, 2001; Moster et al, 2002; Marlow et al, 2005; Lindström et al, 2008; de Vries et al, 2010; van Handel et al, 2010, 2012). Moderate perinatal hypoxia (MPH) may therefore be responsible for less-than-optimal long-term cognitive functioning, which might in turn prevent children to reach their full potential and autonomy. It is thus important to establish experimental animal models to study how MPH changes the developmental trajectory of specific neuronal circuits, thereby contributing to cognitive dysfunctions.

GABAergic circuits comprise an astonishing variety of different cell types, which are likely differentially recruited by distinct behavioral events. An important subtype of GABAergic cells, the fast-spiking, parvalbumin-positive (PV) cells, generate action potentials at high frequency and synchronize the activity of excitatory pyramidal neurons (Hu et al, 2014). PV cells are particularly important for the generation of gamma oscillations, which in turn regulate many cognitive functions including goal-directed attentional processing and working memory (Cardi et al, 2009; Sohal et al, 2009; Buzsáki et al, 2012; Kim et al, 2016). Recent findings indicate that PV cells utilize much more energy than other cortical neurons (Whittaker et al, 2011; Kann et al, 2014), which may render them highly vulnerable to conditions of metabolic and oxidative stress caused by MPH.

The receptor p75NTR is a member of the tumor necrosis factor receptor superfamily, which binds to mature and precursor neurotrophins and regulates numerous functions, including cell death, neuronal differentiation, neurite outgrowth, and synapse pruning. In the mouse nervous system, p75NTR is expressed at high levels in the embryo and in the first postnatal weeks, coinciding with programmed cell death and axonal pruning (Singh et al, 2008; Naska et al, 2010). While p75NTR is generally low in the adult brain, several studies showed that it is upregulated after injury or with disease, such as Alzheimer’s disease, corticospinal axotomy, seizures, hypo-osmolar stress, oxygen–glucose deprivation in organotypic hippocampal slices or focal cerebral ischemia in adult mice *in vivo* (Park et al, 2000; Fahnerstock et al, 2001; Jover et al, 2002; Volosin et al, 2007; Ramos et al, 2007; Irmady et al, 2014). By affecting p75NTR expression, MPH could alter cortical circuit development, thus affecting cognitive function in adult mice.

We established a mouse model of MPH that shows long-term deficits in brain activity and cognition, in parallel to reduced PV expression levels, suggesting that PV cells may be in a more immature state. We further found that the expression level of the neurotrophin receptor p75NTR, which limits PV cell maturation during the first postnatal weeks (Baho et al, 2019), is increased in MPH mice. Embryonic deletion of p75NTR restricted to neurons expressing the transcription factor Nkx2.1, which include cortical PV cells, protects mice from cortical PV expression loss and the long-term cognitive effects of MPH. Finally, adult mice who were treated for one week with a p75NTR inhibitor starting 4 hours after MPH show normal cognitive behavior and gamma oscillation during active exploration as adult mice, suggesting that p75NTR upregulation following MPH contribute to the development of abnormal cortical oscillations and cognitive impairments in adulthood.

## Material and Methods

### Animals

*Ngfr1* (codying for p75NTR) floxed mice with loxP sites flanking exons 4 to 6 of *Ngfr1* gene (*p75NTR^flox/flox^*) were kindly provided by Dr Vesa Kaartinen (Bogenmann et al, 2011). The driver mouse line expressing Cre recombinase, *Nkx2.1-Cre* (Jackson Laboratories, RRID:IMRS_JAX:008661), was crossed to the *Ngfr1* floxed mice and the respective progenies were backcrossed to generate the heterozygous and control genotypes within the same litter. To control for the pattern of expression of Cre, we introduced the RCE allele using Gt(ROSA)26Sortm1.1(CAG-EGFP)Fsh/J mice (Jackson Laboratories; RRID:MMRRC_032037-JAX). The RCE line carries a loxP-flanked STOP cassette upstream of eGFP sequence within the Rosa26 locus. Removal of the loxP-flanked STOP cassette by Cre-mediated recombination allows promoter-specific downstream eGFP expression. All mice were housed under standard pathogen-free conditions in a 12h light/dark cycle with *ad libitum* access to sterilized laboratory chow diet. Animals were treated in accordance with Canadian Council for Animal Care and protocols were approved by the Animal Care Committee of CHU Ste-Justine Research Center.

### Moderate Perinatal Hypoxia (MPH) model

The protocol for moderate perinatal hypoxia (MPH) was adapted from a previously published model by the Jensen lab (Jensen et al, 1991; Sun et al, 2016). Mice pups of both sexes were taken from the dams on either P8 or P9. They were then exposed directly to 4-6% O2 for 10 minutes. This reliably induces seizures with a 5-minute latency, which corresponds clinically to the moderate (stage 2) encephalopathy, a prominent neonatal hallmark defined by the Sarnat Scale (Volpe, 2008). Seizures were identified by observation of motor behavior, in particular mice showed focal ictal manifestations such as arrest of movement (freezing) associated with head bobbing, repetitive limbs movement, and culminating, in most cases, to a generalized tonic–clonic seizure with loss of the righting reflex. Seizures were confirmed by in vivo EEG recording in 2 mice to initially establish the protocol (Suppl. Figure 1B). Age– and sex-matched naïve littermates were taken from their dams at the same time, placed in a separate cage, and given back to the mother at the same time as the pups exposed to MPH.

### Immunohistochemistry

Mice were perfused transcardially with saline followed by 4% Paraformaldehyde (PFA 4%) in phosphate buffer (PB 0.1M, pH 7.2). Brains were post-fixed with 4% PFA overnight and subsequently transferred to a 30% sucrose solution in sodium phosphate-buffer (PBS) for 48hrs. They were then frozen in molds filled with Tissue Tek using a 2-Methylpentane bath cooled with a mixture of dry ice and ethanol (∼ –70°C). Coronal sections of 40 µm were obtained at optimal cutting temperature using a cryostat (Leica Microsystems VT100). Brain sections were blocked in 10% normal goat serum (NGS) and 1% Triton X-100 for 2 hr at RT. Slices were then incubated for 48h at 4°C with the following primary antibodies: rabbit anti-PV (1:5000, Swant, Cat# PV25.RRID:AB_10000344), chicken anti-GFP (1:1000, Abcam, Cat# 13970, RRID:AB_300798); mouse anti-NeuN (mouse monoclonal, 1:400, Millipore; catalog #MAB377, RRID:AB_2298772). This step was followed by incubation with secondary antibodies for 2h at RT to visualize primary antibodies. The secondary antibodies used were Alexa-Fluor conjugated 488, 555, and 647 (1:400, Life technologies; 1:1000, Cell Signaling Technology). After rinsing in PBS (three times), the slices were mounted in Vectashield mounting medium (Vector).

### RNAscope

Fluorescent multiplex RNAscope were performed as described in (Baho et al, 2019). Non-perfused frozen brain sections (15μm) were cut using a cryostat (Leica Microsystems VT100) and mounted on Superfrost Plus Gold Glass Slides (Fisher Scientific, #22–035-813). Slides were subsequently stored at –80°C. Probes against Ngfr mRNA (494261), which codes for p75NTR, and Pvalb mRNA (421931-C2), which codes for PV, as well as all other reagents for ISH and DAPI labeling, were purchased from Advanced Cell Diagnostics. The tissue pretreatment, hybridization, amplification, and detection were performed according to User manual for fixed frozen tissue (Advanced Cell Diagnostics). During RNAscope hybridization, positive probes (catalog #320881), negative probes (catalog #320871), and Pvalb/Ngfr probes were processed simultaneously. Briefly, the slides were removed from –80°C and rinsed with PBS. They were submerged into 1 Target retrieval solution (catalog #322000) for 5 min at 100°C, and then rinsed in distilled water followed by 100% EtOH dip to remove access water. Protease III (catalog #322337) was added to each section and incubated for 30 min at 40°C followed by washing in distilled water. For detection, probes were added to each section and incubated for 2 h at 40°C. Unbound probes were subsequently washed away by rinsing slides in 1 wash buffer (catalog #310091). AMP reagents (AMP1 catalog #320852, AMP2 catalog #320853, AMP3 catalog #320854, AMP4A catalog #320855, C1 probes–Alexa-488; C2 probes-Atto-550) were added to each section and incubated for as per the manufacturer’s instructions, and washed in wash buffer for 2 min. Sections were stained with DAPI (catalog #320858) for 30s, and then mounted with Prolong Gold Antifade Mountant (Invitrogen, catalog #P36961). This experiment was performed using tissue from mice from 3 different litters. Half of the litters was exposed to hypoxia at P9, while half of the littermates were used as control (sham, separated from the dam for the same amount of time). Tissue was harvested 48h following MPH.

### Confocal imaging and analysis

For each data acquisition and analysis, the experimenter was blind to treatment (naïve vs MPH) until the completion of data analysis. <u>RNAscope</u> analysis was performed as described in Lavertu-Jolin et al (2023). Briefly, images were acquired with a Leica Microsystems confocal microscope (SP8-STED), using an 63×/1.4 oil objective, and deconvolved using Huygen’s express deconvolution option. The figures represent the maximum projection of acquired z stack. To determine the number of *Ngfr1* or *Pvalb* dots per *Pvalb*+ cell, images were analyzed using a custom-made ImageJ-Fiji macro. Briefly, a dark region was selected on each focal plane to measure the mean gray value (MGV) as background. Then, for each channel and each focal plane, the background value was subtracted, and intensity value of each pixel was readjusted by multiplying it with 65535 (65535-background). Then, each *Pvalb*+ cell was manually selected based on the presence of *Pvalb* puncta, filtered with the Gaussian Blur 3D option and analyzed using the 3D objects counter (threshold= 4500, minimal voxel size= 6). Data are normalized to the mean mRNA puncta (number of object) detected in naïve mice in each experiment.

<u>Immunohistochemistry</u> analysis of PV sections were processed in parallel, and all images were acquired the same day using identical confocal parameters. Confocal images (Leica Microsystems, SP8 confocal microscope) were acquired using either a 20× water-immersion objective (NA 0.7; Leica Microsystems) or a 63× glycerol objective (NA 1.3; Leica Microsystems). For each animal, we acquired at least 3 confocal stacks in layers 3-5 in both hemispheres. Data were obtained from 3 or 4 brain sections per animal. z stacks were acquired with a 1μm step, exported as TIFF files, and analyzed using ImageJ software (RRID:SCR_003070). Briefly, PV rings (between 7 and 10 in each stack) were outlined, and the mean gray values were measured, after background subtraction. Background was determined by measuring mean gray values in at least three different areas (ROI), where immunolabeling was absent, in the same focal plane where PV perisomatic rings were selected. The number of PV cells or GFP-expressing cells was analysed using Neurolucida (MicroBrightField) software.

### Western Blot

Western blots were performed as described in (Baho et al, 2019). Cortical tissue extracts were quickly frozen in nitrogen and stored at –80°C until protein extraction procedure. Samples were dissociated in lysis buffer (2 mM EDTA, 1% Igepal C-630 in TBS, 50 mM Tris, 150mM NaCl, pH 7.6) containing protease inhibitor (Roche Diagnostics, catalog #11836153001) and phosphatase inhibitor (Roche Diagnostics, catalog #04906845001) mixtures at 4°C, centrifuged 10 min at 10,000 x g at 4°C, and the supernatants were dosed with Bradford buffer (Bio-Rad, catalog #5000006). All samples used for Western blot analysis of p75NTR expression were run on the same gel. Samples were diluted at the same concentration in Laemmli solution (20% glycerol, 4% SDS, 10% 2,6-mercaptoethanol, 0.02% bromophenol blue in 125mM Tris, pH 6.8) and boiled at 95°C for 5 min; 20μg of protein was migrated on precast gel, 4%–15% acrylamide (Bio-Rad, catalog #456 –1086) at 185V for 40 min. The proteins were transferred to a PVDF membrane (Millipore, catalog #IPVH00010) at 100V for 30 min in transfer buffer (20% methanol, 192 mM glycine in 25mM Tris). The membranes were blocked in 5% blocking solution (Bio-Rad, catalog #170–6404) in TBS/T during 2h at room temperature. Membranes were then probed with anti-p75NTR (1:200; rabbit monoclonal, kindly gifted by P. Barker) and anti-GAPDH 1:8000 (mouse monoclonal IgG; Thermo Fisher Scientific, catalog #AM4300, RRID:AB_2536381) in 5% blocking solution/TBST (0.1% Tween in TBS) overnight at 4°C. The membranes were washed in TBST (3×15 min at room temperature) and probed with the following secondary antibodies, anti-mouse-HRP (1:6500, Sigma-Aldrich catalog #A4416, RRID:AB_258167) and anti-rabbit-HRP (1:10,000, Abcam, catalog #ab6721, RRID:AB_955447), for 2h at room temperature. The membranes were washed in TBST (3×15 min) and revealed with ECl (PerkinElmer, catalog #NEL_103001EA). Membranes were exposed to Bioflex MSI autoradiography/x-ray film for different time intervals, and only the films that showed easily identifiable, but not saturated, bands for every sample were used for quantification, using ImageJ software (RRID:SCR_003070; http://imagej.nih. gov/ij). Background mean gray value was subtracted, and then values were normalized on GAPDH mean gray value. Specificity of the anti-p75NTR antibody was verified using brain lysates from p75NTR KO mice. The experimenter was blind to treatment (naïve vs MPH) until the end of data analysis.

### Drug treatment

Mice pups were administered the p75NTR antagonist LM11A-31 via gavage (50mg/kg) or its vehicle solution (saline), starting 4h after MPH exposure at P8/9 and then twice per day for the following 7 days, until the mouse pups were P15/16. LM11A-31 was custom manufactured, purified and its structure and purity confirmed as previously described (Yang et al, 2020).

### Behavioral tests

All tests were performed using mice of both sexes aged between P80-P120. The experimenter was blind to treatment (naïve vs MPH) and drug treatment until the end of data analysis.

#### Elevated plus maze (EPM)

The apparatus consisted of two open arms without walls across from each other and perpendicular to two closed arms with walls joining at a central platform. The full apparatus is elevated above the floor. Each mouse was placed at the junction of the two open and closed arms and allowed to explore the maze during 5 min while video recorded. The recorded video file was analyzed with the SMART video tracking system (v3.0, Harvard Apparatus) to evaluate the percentage of time spent in the open arms (time spend in open arms / (total time spent in open + closed arms) x 100).

#### Open Field (OF)

Each subject mouse was gently placed at the center of the open-field arena (44.60cm x 44.60cm) and recorded by a video camera for 10 min. The recorded video file was analyzed with the SMART video tracking system (v3.0, Harvard Apparatus). The open field arena was cleaned with 70% ethanol and wiped with paper towels between each trial. We analyzed the time spent in the center (45% of the surface) versus the periphery and the total distance traveled.

#### Object location recognition (OLR) and novel object recognition (NOR)

The previously mentioned open field box was utilized as the testing chamber for the OLR and NOR procedure. The test was performed under dim lighting at 12 lux in a quiet environment and recorded by SMART video tracking system (v3.0, Harvard Apparatus). This procedure consisted of 7 steps. 1) Habituation stage: the mouse was placed in the testing chamber and allowed to freely explore for 5 minutes; followed by 2) 1h interval in a clean waiting cage. 3) Familiar stage: 2 identical objects were placed approximately 30 cm away from the starting wall and 15 cm away from the relative side walls. The mouse was first placed at the middle of the starting wall while facing it and then allowed to freely explore for 10 minutes. This step was followed by 4) 1h in the waiting cage. 5) OLR stage: one of the familiar objects was moved to 15 cm from the starting wall, thus becoming the displaced object. The mouse was placed in the same initial place as for the familiar stage and allowed to explore for 10 minutes. 6) This was again followed by 1h wait in the waiting cage. 7) NOR stage: The familiar object that was not displaced previously was then replaced by a novel object at the same spot. The mouse was then allowed to explore freely for 10 minutes. Discrimination Index = (exploration time (s) with novel object – exploration time (s) with familiar object) / (exploration time (s) with novel object + exploration time (s) with familiar object). Only mice that explored the objects for a minimum of 5 seconds were included in the analysis.

#### Attention set shifting task (ASST)

This test was performed as described in Chehrazi et al, 2023. Briefly, behavioral experiments were performed in a home-made acrylic rectangular-shaped maze (30 cm long × 20 cm wide × 18 cm high) divided in half by a guillotine-like door that extended through the width of the maze. One half served as the start area and the other half served as the choice area. In the choice area, an acrylic wall (12 cm long × 18 cm high) presenting four ¼” diameter sniffing holes on the bottom extended out from the back wall and divided the choice area into two equally sized and distinct spatial locations. Each of these two locations contained a ceramic ramekin identical in color and size (non-porous; depth 3.5 cm × diameter 6 cm). A ramekin filled with water was placed in the starting area.

A piece of Cheerios (approximately 20 mg in weight) was used as the food reward. The cues, either olfactory (odor) or somatosensory and visual (texture of the digging medium which hides the food reward), were altered and counterbalanced. All odors were essential oils, and unscented digging media. The testing pots were scented a day before to allow the scent to dissipate by adding approximately 0.1 ml of essential oil to the top of the filter paper using a syringe with a 25G needle.

The experiment was divided into five steps: handling, food restriction, acclimation, training, and testing as explained in Heisler et al (2015). First, mice were single-housed and habituated to a reverse light/dark cycle for at least one week. Mice doing this test were food restricted to 85% of their ad lib. feeding weight in the days prior to testing and for the full duration of the experiment. All mice were handled 2-3 min per day starting 8 days prior to the first day of food restriction and their body weights were recorded each day of handling.

Mice were tested in seven stages: Simple discrimination (SD), Compound Discrimination (CD), Reversal (Rev), Intra-Dimensional Shift 1-3 (IDS1, IDS2, IDS3) and Extra-dimensional shift (EDS). Following SD, media and odor were paired for each of the steps. Rewarding ramekin was placed randomly on the left or right side of the maze to avoid the use of spatial cues. Food reward was covered with scented (or not, in the case of SD) media and cereal dust was sprinkled on top of the media to avoid aberrant smell cues. To complete a trial, the mice had to dig into a media to retrieve the food reward. In incorrect trials, animals were free to visit the other side of the apparatus, but the other ramekin was removed. At the end of each trial, animals were guided to the start area and the ramekins were refilled or changed. The testing phase was scheduled on 2 days. SD, CD, CDR and IDS1 were performed on day 1, followed by IDS2, IDS3 and EDS on day 2.

Odor, medium combination and location (left or right) of the baited bowl was determined as in Chehrazi et al, 2023. At each stage of the task, while the particular odor-medium combination present in each of the two bowls may have changed, the particular stimulus that signaled the presence of food reward remained constant over each portion of the task (initial association, rule shift, and rule reversal). In our case, the initial association was paired with a specific digging medium with food reward, and the odor was considered the irrelevant dimension. The mouse was considered to have learned the initial association between stimulus and reward if it made 8 correct choices in 50 consecutive trials. Each portion of the task ended when the mouse met this criterion. If a mouse did not make 8 correct choices consecutively within 50 trials, it failed that stage (indicated as an X on the graph) and was not tested on the subsequent steps. 6 consecutive no choices (3 min without a choice) were considered as a failure to participate and the mouse was moved from the experiment.

Following the initial association, the rule-reversal portion of the task began, during which the stimulus that had been consistently not rewarded during the initial association becomes associated with reward. For example, if paper had been associated with reward during the initial association, then during the rule reversal, straw would become associated with reward. Finally, during the extra-dimensional shift stage, the irrelevant dimension (odor in our case) became the relevant dimension.

### Video-EEG recordings for ictal activity detection

To analyze the occurrence of ictal activity in our model, we monitored adult MPH and naïve mice by continuous video-EEG recordings after implantation of electrodes in the hippocampus and the overlying cortex. The insertion of electrodes was performed under isoflurane anesthesia, the head of the mouse being immobilized in a stereotaxic frame. A stainless-steel bipolar electrode (Plastics-1 Inc., Roanoke, VA, USA) one cut to 1.5mm depth (hippocampal electrode) and the other at 0.8mm (cortical electrode) was positioned at the following coordinates with reference to the Bregma: AP −1.5 (posterior to Bregma), ML 2.0 lateral to midline. The reference electrode was placed in the cerebellum. Following a 10-day recovery period, EEGs were recorded with a Stellate systems 32-channel video-EEG recording unit, beginning at P60 till P100. Mice were monitored twice/24h for two 4-hour sessions (day session and night session) over a 4 weeks period. Two separate observers blind to the treatment groups analyzed the recordings. Further, during the initial establishment of the protocol, we implanted 2 mouse pups at P7, two days before exposing them to the MPH protocol at P9 and monitored their EEG activity to confirm seizure occurrence during hypoxia exposure.

### *In vivo* brain oscillation recordings

#### Surgery

Cranial surgery was performed under deep anaesthesia, induced by an intraperitoneal injection of ketamine/xylazine cocktail. A craniotomy was performed at the coordinates described below on the right side of the brains and custom-made clusters of four to six 50 mm-diameter-insulated-tungsten wires were implanted. In each cluster, wires tips were 50–150 mm apart. A custom-designed and 3D-printed microdrive-like scaffold held the three electrode clusters at fixed positions, thus allowing the simultaneous implantation of all three and reducing surgery time. Data were acquired with a custom-made headstage using Intan Technologies’ RHD Electrophysiology Amplifier chip. A ground and reference wire was also gently introduced in the contralateral frontal lobe and a 2cm carbon-fibre bar was placed on the back of the skull. The microdrive apparatus, the carbon-fibre bar and the ground-and-reference wire were then daubed with dental acrylic to encase the electrode-microdrive assembly and anchor it to the skull. For 1 week after surgery, animals were treated with Metacam and the skin around the implant was disinfected daily.

Electrode clusters were implanted in 4 different cortical regions at the following coordinates: prelimbic cortex (PL), anteroposterior +2.00mm, mediolateral +0.25mm from bregma and 1.5mm deep; primary auditory cortex (A1), –2.5 mm posterior to bregma and 3.9 mm lateral to the midline posterior parietal cortex, 0.8 mm deep; primary motor cortex (M1), –1.5 mm posterior to bregma and 1.5 mm lateral to the midline. This cluster targeted both the supra and infragranular layers and also included two longer (1.2 and 1.3mm) wires reaching the dorsal region of the hippocampus (CA1 pyramidal layer and radiatum). The correct depth in the CA1 was confirmed by the presence of ripples, which appeared on the recordings when the electrode reached the pyramidal layer. For this study, we analysed only signals from cortical layer 5 and the hippocampal radiatum layer.

#### Data acquisition

One week after surgery, animals were introduced for 30 minutes in novel open field. We used a dimly illuminated open field environment (45×35cm), surrounded by 60cm-high transparent walls, and equipped with video monitoring. The walls were sprayed and wiped clean with 70% ethanol 30 minutes before the introduction of each animal. Video and EEG signal were simultaneously acquired. Video sampling rate was 15 frames per second and EEG sampling rate was 20 KHz. EEG recordings were performed using an open-ephys GUI platform (https://open-ephys.org/).

### Data Analysis

#### Brain state classification and data segmentation

Periods of active exploration can be detected by brain state classification. Using the spectral properties of the hippocampal LFP signal we can find epochs of higher theta power which are well known to be related to high arousal states, such as exploring, sniffing, and whisking. In contrast, periods of non-locomotor behaviours (or low arousal states) such as immobility, eating, drinking, and grooming, are characterized by large irregular activity oscillating at delta frequency. Here, active exploration and non-explorative periods were detected using the ratio of power in theta (6–11 Hz) range to power in delta (1–4 Hz) frequency band (Csicsvari J et al, 1999; Malezieux et al, 2020). To extract the power of both frequency bands over time, we used the chronux toolbox function *mtspecgramc* to calculate the multitaper spectrum using a one-second moving window and 0.05 second step. We then performed the theta–delta power ratio and automatically marked periods of theta or delta activity based on whether the threshold detection was higher (theta) or lower (delta) than the ratio at each time point. This threshold was defined as:

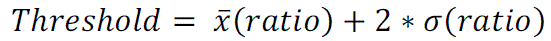

Only periods longer than 2 seconds were considered for analysis. Both, exploratory and non-exploratory periods were then segmented and sequentially organized in a matrix of 2 seconds subperiods.

#### Power Spectrum Density

Power Spectrum Density (PSD) was calculated over both, exploratory and non-exploratory periods using multitaper methods in order to increase spectral resolution. In this case, another chronux function called *mtspectrumc* was used to perform this analysis. Because we were interested in investigating evoked gamma power during active exploration, we performed a baseline correction using non-exploratory periods PSD as baseline, following the formula:

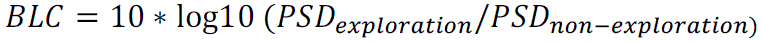

#### Phase Amplitude Coupling

Phase amplitude coupling (PAC) analysis was performed as described in Carren-Munoz et al, 2022. PAC analysis allows the evaluation of nesting oscillations, where the phase of the lower frequency (here called fp) modulates the amplitude of the higher frequency (fa). The bandwidth of the filter used to obtain ‘fp’ and ‘fa’ is a crucial parameter in calculating PAC. The filters for extracting ‘fa’ need to be wide enough to capture the centre frequency ± the modulating ‘fp’. If this condition is not met, then PAC may not be detected even if present. We therefore decided to use a wide enough bandwidth for both ‘fa’ and ‘fb’. The coupling between ‘fp’ and ‘fa’ was quantified using the mean vector length modulation index (MI) as it has been suggested to be more sensitive to coupling strength and width compared with other methods. This approach estimates PAC from a signal with length N, by combining phase (/) and amplitude information to create a complex-valued signal.

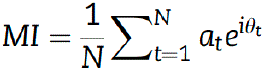

where N is the total number of data-points, t is each time point, at is the amplitude of ‘fa’ at the time point t and ht is the phase angle of ‘fp’ at time point t. To evaluate whether the observed modulation index actually differs from what would be expected by chance, surrogate analysis needs to be performed. To do so, surrogate data are created by shuffling trial and phase-carrying information (500 surrogates) to normalize MI-values. These random permutations create a new time series with broken temporal relationships between the phase and amplitude information. Then the MI is estimated again using the shuffled time series to obtain the null distribution of surrogate modulation index values. A normalized MI (MInorm) is then obtained as a z-score:

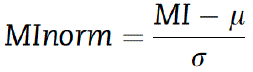

where l and r are the mean and standard deviation obtained from the null distribution. A Bonferroni corrected t-test was then performed and all MI-values exceeding a 5% significance threshold were grouped into clusters for further statistical analysis by group.

### Statistics and reproducibility

Statistical analyses were performed using Prism 9.0 (GraphPad Software). Prior to making comparisons across values, the normality of distribution was tested. Differences between 2 experimental groups were assessed using two-tailed t-test or t-test with Welch’s correction for normally distributed data and Mann Whitney test for not normally distributed data. Differences between >2 experimental groups were assessed using One-Way ANOVA for normally distributed data and One-Way ANOVA on Ranks for not normally distributed data. All bar graphs represent mean ± SEM.

## Results

### Mice that experienced moderate perinatal hypoxia show cognitive impairment in adulthood

Determining how brain development correlates between humans and rodents is not straightforward, however there is also considerable cross-species alignment in terms of key developmental milestones. Based on biochemical and neuroanatomical changes during early development, the general consensus is that a P8-10 rodent brain corresponds roughly to the brain of a term infant (Semple et al, 2013). Therefore, we used this time window as reference to develop a preclinical mouse model of MPH. We first established a protocol that allows us to reliably observe hypoxia-induced seizures in postnatal mice. We found that exposing P8-9 pups directly to 4-6% O_2_ for 8 minutes reliably induces myoclonic and electrographic seizures with a latency of about 5min in 3 separate mouse strains (FVB, C57Bl/6, 129S6; Suppl. Figure 1). This aspect is clinically relevant as seizures are the most prominent neonatal hallmark of Stage 2 (Moderate) encephalopathy as defined by the Sarnat Scale (Volpe, 2008). About 50% of children experiencing hypoxia-induced seizures do not develop identifiable brain lesions or serious motor problems (Gluckman et al, 2005; Shankaran et al, 2005) as adults. On the other hand, recent data suggest that this patient population may be more prone to the development of subtle but more functionally challenging cognitive deficits than previously thought (Dilenge et al, 2001; Maneru et al, 2001; Moster et al, 2002; Marlow et al, 2005; Lindström et al, 2008; de Vries et al, 2010; van Handel et al, 2010, 2012). Thus, as a next step, we sought to characterize the long-term consequences of MPH on brain function, by testing whether MPH mice developed behavioral deficits in adulthood. We found no difference in the open field test, which rules out the presence of major motor problems, and in the elevated-plus maze, which indicates normal anxiety levels (Fig. 1B1, 2; C). However, in contrast to naïve mice, MPH mice did not spend more time exploring a novel object or an object that was displaced compared to a familiar or stationary one (Fig.1E1,2; F1,2), suggesting the presence of recognition and spatial memory deficits. Further, we used the attentional set shifting task (Fig.2B1,2), which is equivalent to the Wisconsin Card Sorting Task (WCST) used in humans to assess prefrontal cortex mediated cognitive flexibility. During this test (Fig.2A), the mice were presented with two bowls on each trial and had to choose to dig in one bowl to find the food reward. Each bowl contained a different odor and a different digging medium, and the odor-medium combinations varied from trial to trial (Fig.2A). During the initial association phase, the food restricted mouse learnt to associate the relevant cue (a specific digging medium) and ignore an irrelevant cue (odor), by pairing a food reward with the medium (initial association, simple discrimination SD, followed by compound discrimination CD). During the reversal stage, the mouse had to reverse rules and learn that the food was now associated with the digging medium that had not been rewarded during the initial association (Rev1). The stimulus-reward association was then reinforced in subsequent tasks where the type of digging medium and odor changed, but the paired association between medium and reward remained (intra-dimensional shift, IDS1, IDS2, IDS3). During the extra-dimensional shift stage, the irrelevant dimension (odor) became the relevant dimension (EDS). We observed that while both control and MPH mice were able to learn the initial association in a similar number of trials, MPH mice performed significantly worse in the subsequent steps, since they were unable to modify their response when the rules changed during different stages of the test (Fig.2B1,2).

**Figure 1.**
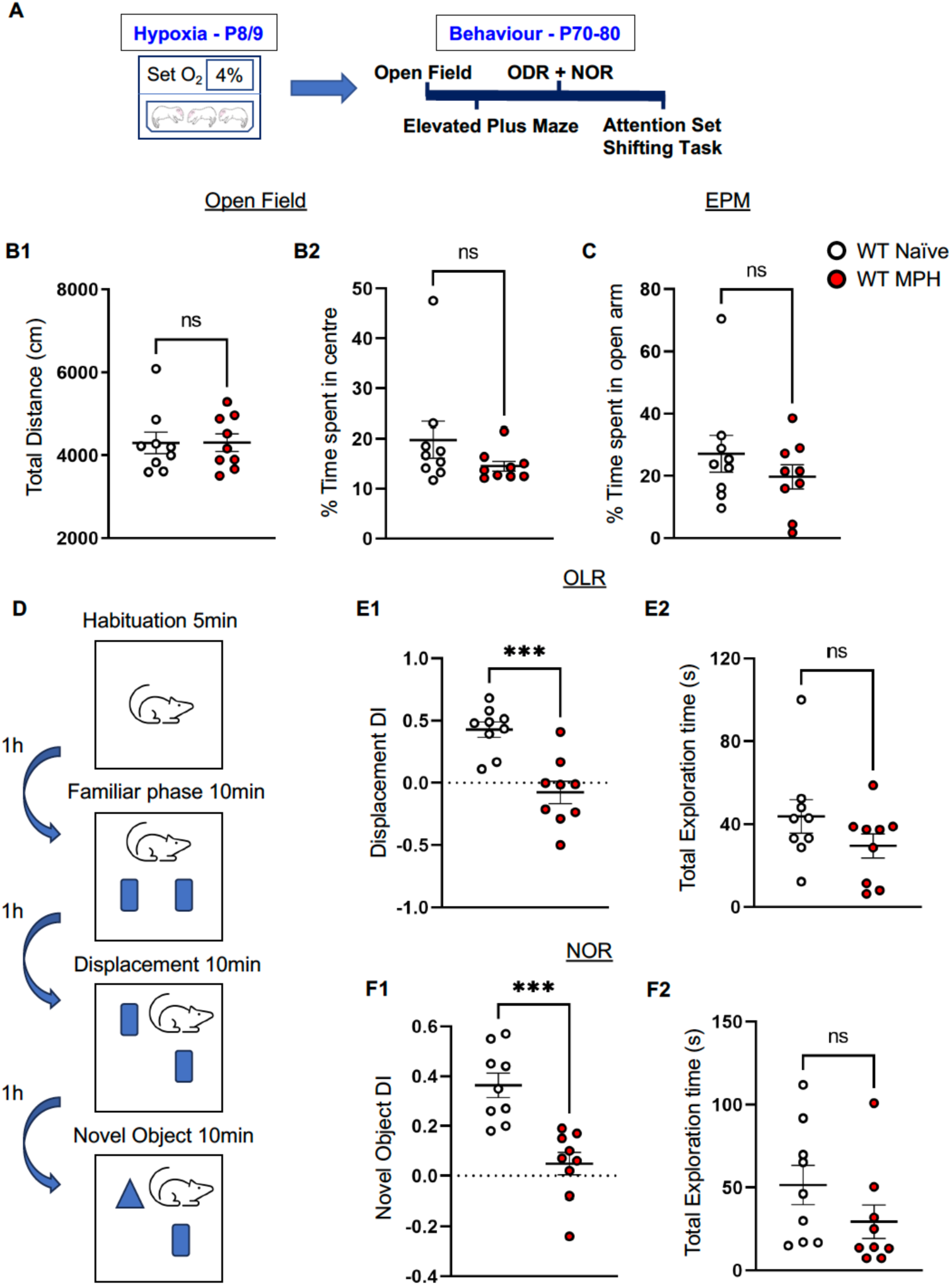
Adult MPH mice show cognitive dysfunctions in object recognition and spatial memory. **A.** Behavioural testing protocol in adult mice following moderate perinatal hypoxia (MPH). **B1,2.** MPH and naïve mice do not show any significant difference in total distance covered (Mann-Whitney test (MW) U=38; p=0.8633) or time spent in the center of the open field (OF, MW U=23; p=0.1359). **C.** MPH and naïve mice do not show any significant differences in the time spent in closed arms of the elevated plus maze (EPM, MW U=32; p=0.4894). **D**. Schema of OLR and NOR behavioural protocol. **E, F.** MPH mice show deficits in spatial (E1, OLR, MW U=4; p=0.0005) and novel object recognition memory (F1, NOR, MW U=1; p<0.0001), but no difference in both OLR and NOR total exploration time (E2, OLR total, MW U= 25, p=0.1903; F2, NOR total, MW U=20, p=0.0770). **B, C, E, F,** Naïve n=9 and MPH n=9 mice; Lines represent Mean ± SEM, while circles represent values for a single mouse.

**Figure 2.**
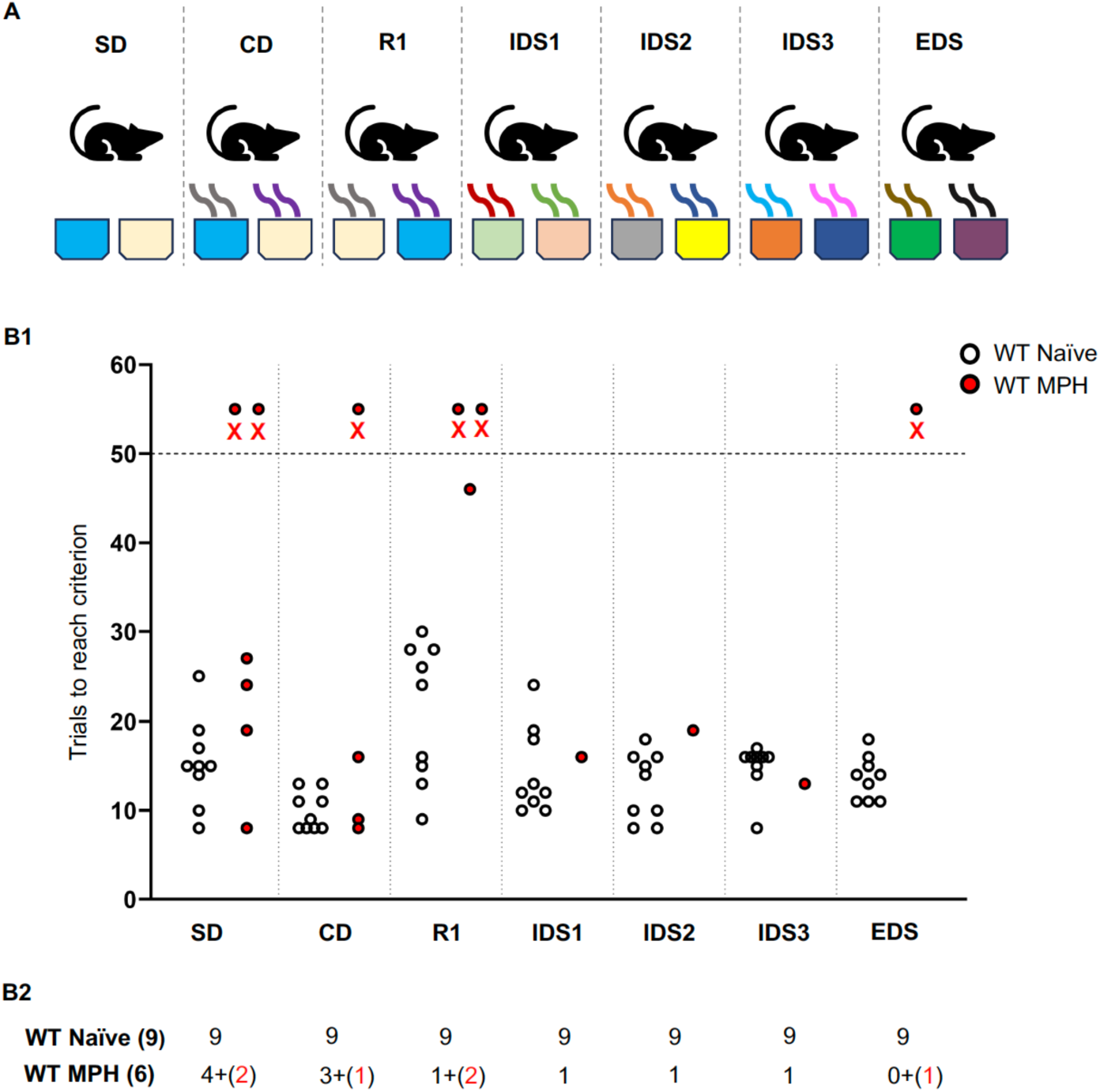
Adult MPH mice show deficits in cognitive flexibility. **A.** Schema showing the steps in the ASST behavioural task. **B1.** MPH WT mice fail to complete all the steps of the ASST, red crosses show the mice that fail at any given step, (not reaching criterion within 50 trials), as compared to the Naïve WT controls. **B2.** Table summarizing the overall results of ASST for Naïve vs MPH mice. The number of mice that failed to pass each indicated stage are in red, while those that pass it are in black. **B1, B2** Naïve n=9 and MPH n=6 mice, circles represent values for a single mouse.

Previous work showed that perinatal rats exposed to a similar hypoxia protocol developed spontaneous seizure as adults (Rakhade et al, 2011). Since behavioral performance may be affected by spontaneous seizures, we used video-EEG to detect the occurrence of spontaneous motor and electrographic seizures. Adult mice (5 naïve and 6 MPH) were monitored twice/24h for two 4-hour sessions (day session and night session) over a 4 weeks period. Over this period, we did not detect seizures in either experimental group, suggesting that mice exposed to MPH do not develop spontaneous seizures as adults (Data not shown).

Altogether, the behavioral analysis supports the conclusion that mice which experience moderate perinatal hypoxia develop cognitive impairment in the memory and cognitive flexibility domains.

### p75NTR expression is increased 48h after MPH particularly in PV cells

Several lines of evidence suggest that increased p75NTR expression occurs after brain insults and may play a major role following brain injury. In particular, Irmady and collaborators (2014) showed that p75NTR expression was rapidly increased after *in vivo* focal cerebral ischemia in adult mice or after oxygen–glucose deprivation in organotypic hippocampal slices. In addition, apoptosis following oxygen–glucose deprivation was reduced either in hippocampal slices from wild-type mice treated with function-blocking antibodies to p75NTR or in hippocampal slices from p75NTR^+/-^ (haploinsufficient) mice. Finally, p75NTR KO mice showed a significant decrease in infarct volume compared to wild type mice following focal cerebral ischemia (Irmady et al, 2014). We thus sought to explore whether p75NTR expression was affected by hypoxia in perinatal mice. Western blot analysis showed significantly increased p75NTR expression in cortical lysates from MPH mice compared to control littermates 48 hours after hypoxia exposure (Fig.3A2,B; Suppl Figure 2A). Changes in p75NTR expression may occur in different neuronal populations and even in glial cells. To explore whether p75NTR upregulation occurred in specific cell types, we used RNAscope multiplex fluorescent ISH (Advanced Cell Diagnostics), which allows single-mRNA molecule detection, to label simultaneously the mRNAs coding for p75NTR (gene name: *Ngfr)* and PV (gene name: *Pvalb*) in brain slices from mice, which were exposed to MPH 48hrs prior and control littermates (Fig.3C, D). We found a significant increase in the number of *Ngfr1* puncta, but not *Pvalb* puncta, in cortical neurons coexpressing both mRNAs from mice exposed to MPH compared to their controls (Fig.3E,G). Conversely, the number of *Ngfr1* puncta in cortical neurons lacking *Pvalb* expression, which most likely represent pyramidal glutamatergic neurons, was not different in the two experimental groups (Fig.3F). This observation suggests that perinatal hypoxia increases p75NTR expression primarily in PV cells.

**Figure 3.**
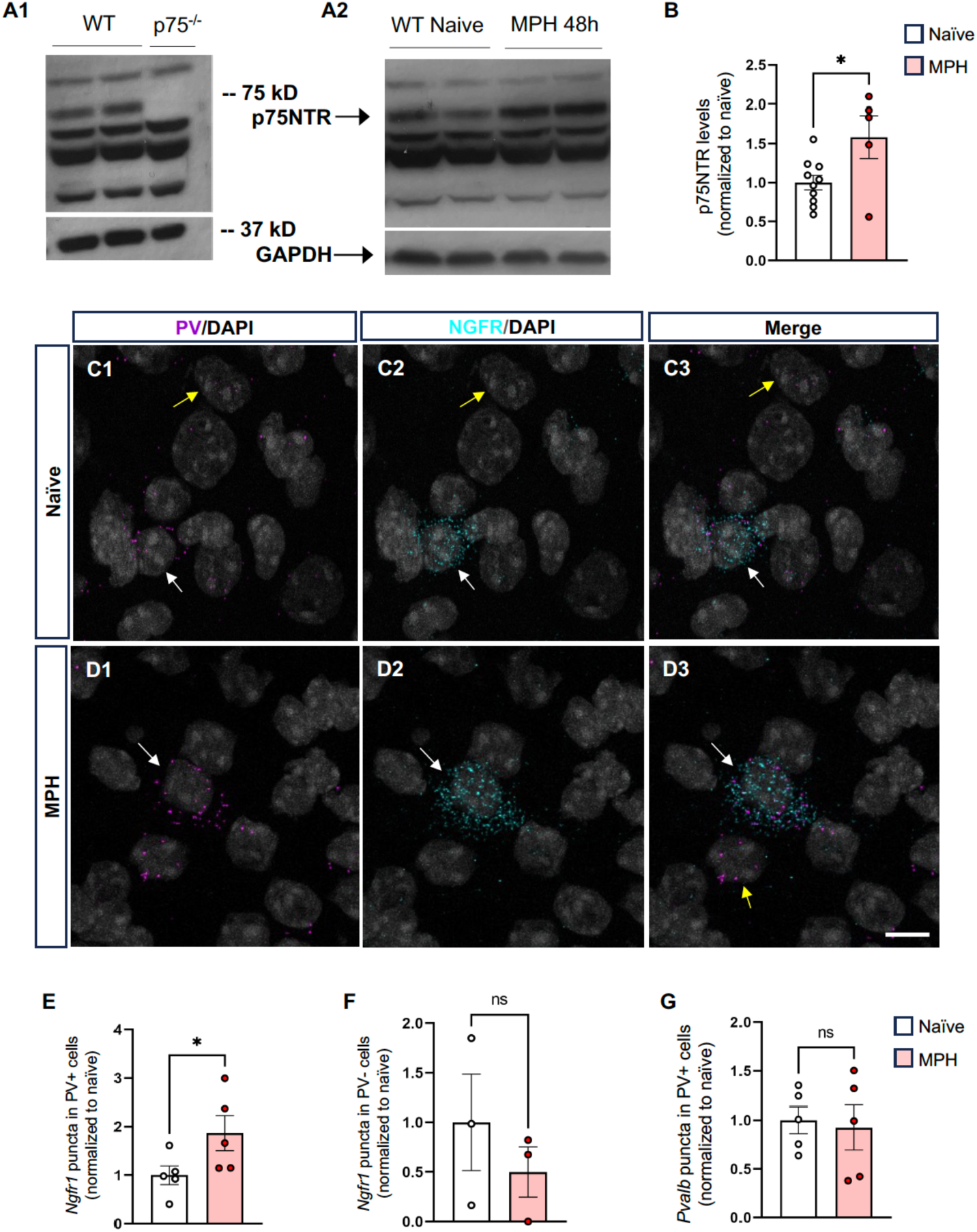
Cortical p75NTR expression is significantly increased 48 hrs after MPH exposure. **A1**. Western blot confirming the specificity of the anti-p75 NTR antibody. Note that the band at 75 kD is absent in cortical lysates from p75NTR knockout (p75NTR^-/-^) mice. **A2, B** Western blot bands and analysis of p75NTR expression levels in cortical lysates of mice exposed to MPH and naïve littermates, 48 hrs after (MW U=11; p=0.04). Different lanes represent different mice. Naïve n=10 and MPH n=5 mice. **C.** p75NTR mRNA levels are significantly increased in PV cells 48 hrs after exposure to MPH. **C1-6**. Representative images of fluorescent RNAscope in situ hybridization against *Ngfr1* (p75NTR, cyan) and *Pvalb* mRNA (PV, magenta) from naïve (C1-3) vs MPH WT mice. Cell nuclei are labeled by DAPI (grey). White arrows point to somata expressing *Pvalb* mRNA, yellow arrows point to non-*Pvalb* mRNA expressing somata. Scale bar: 10 μm. **E.** Normalized number of *Ngfr1* puncta in cells expressing *Pvalb* show significant difference between naïve and MPH mice (MW U=2, p=0.0317). **F**. Normalized number of *Ngfr1* puncta in cells negative for *Pvalb* however show no difference (MW U=2, p=0.4). **G**. Normalized number of *Pvalb* puncta in MPH and naïve mice is not significantly different (MW U=11, p=0.8413). N=5 for both naïve and MPH WT mice.

### p75NTR expression in cortical PV cells render them vulnerable to MPH

PV cells undergo a dramatic change in expression profile around P10-14, when they start to form multiple synapses onto target cells (Chattopadhyaya et al, 2004; Okaty et al, 2009). In particular, down-regulation of p75NTR expression promotes the formation of PV cell synapses (Baho et al, 2019). Thus, alterations in p75NTR levels following MPH may affect PV cell maturation per se.

To address this question, we analysed cortical PV cell density and found that it was significantly reduced in MPH mice compared to control littermates (Fig.4 A,B,E). Further, we found that the intensity PV+ perisomatic puncta, which largely correspond to GABAergic synapses made by PV cells, was significantly reduced in adult MPH mouse cortex (Fig.4 F,G,J).

**Figure 4.**
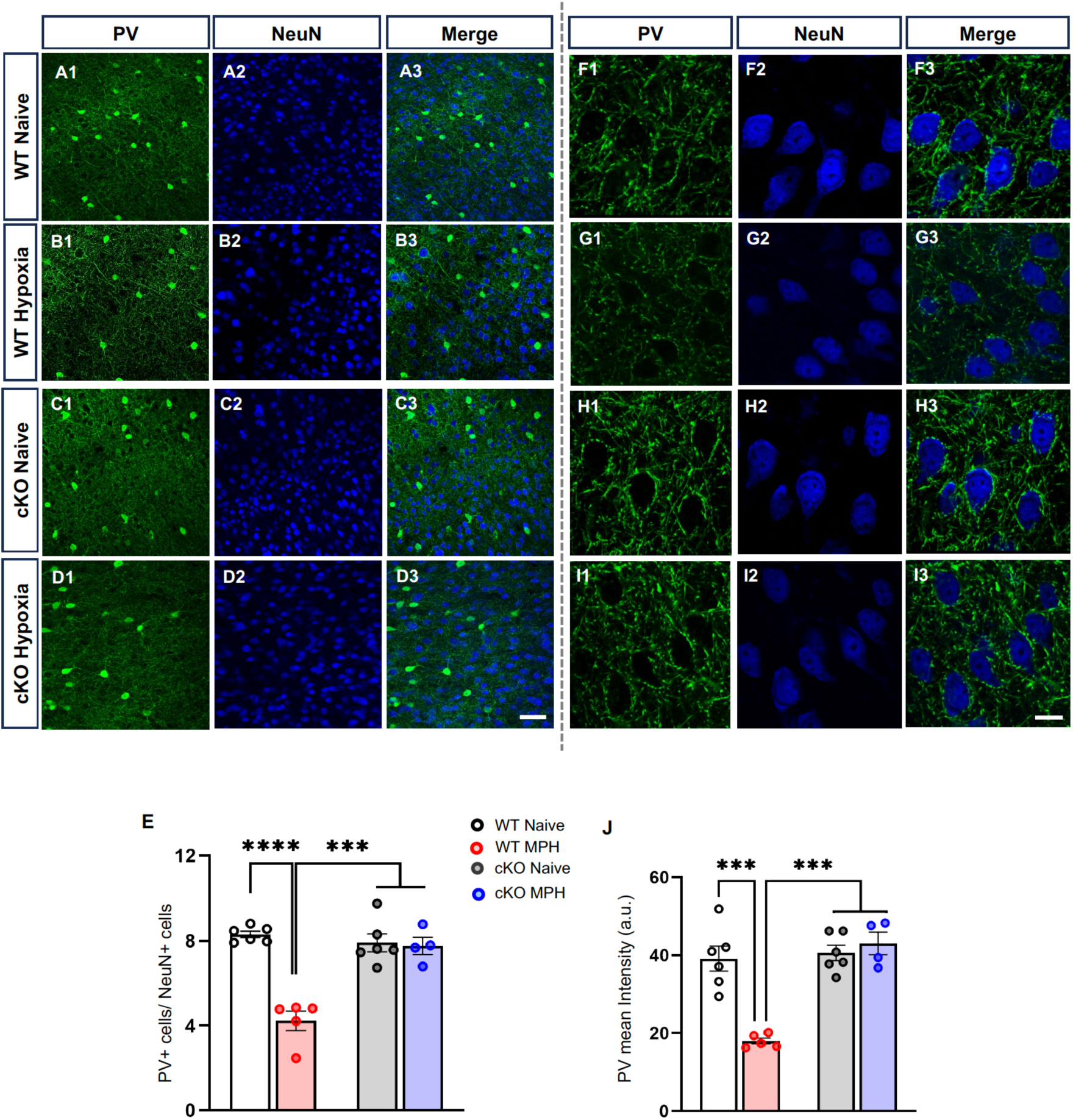
Genetic deletion of p75NTR in MGE-derived GABAergic neurons prevents long-term alterations in PV expression in mice exposed to MPH. **A-D**. Representative images of somatosensory cortex from naïve and MPH exposed (*Nkx2.1-Cre;p75^+/+^*) WT and (*Nkx2.1-Cre;p75^flx/flx^*) cKO littermates immunolabeled for PV (green) and NeuN (blue). Scale bar: 30μm. **B1-3**. The density of cortical PV+ somata is significantly reduced in the somatosensory cortex of adult WT MPH mice compared to WT naïve mice (**A1-3, E**, 2-way ANOVA, Tukey’s *post hoc* multiple comparison test, MPH-WT vs naïve-WT: p<0.0001; MPH-WT vs naive-cKO: p<0.0001; MPH-WT vs MPH-cKO: p=0.0001). On the other hand, the densities of PV+ somata in cKO MPH mice is not significantly different from their naïve control group (**C1-3, D1-3, E**, Tukey’s *post hoc* multiple comparison test, Naive-cKO vs MPH-cKO p>0.9999). **F-I**. Representative high magnification images of perisomatic PV-positive puncta (green) in naïve and MPH WT and cKO mice. Scale bar: 5μm. **J**. The intensity of PV-positive perisomatic puncta around pyramidal neurons, which represent putative synapses made by PV cells, is significantly reduced in WT MPH mice compared to the other 3 experimental groups (2-way ANOVA, Tukey’s *post hoc* multiple comparison test, MPH-WT vs Naïve-WT: p=0.0005; MPH-WT vs Naïve-cKO: p=0.0003; MPH-WT vs MPH-cKO: p=0.0004; as compared to MPH-cKO vs Naïve-cKO: p=0.9906; MPH-cKO vs Naïve-WT: p=0.9169; Naïve-WT vs Naïve-cKO: p=0.9771;). Naïve WT: n=6 and cKO n=6 mice. MPH WT: n= 5 and cKO n=4 mice.

An issue when using PV immunostaining to identify PV cells is that PV expression starts late during development (after the first postnatal week) and its expression level depends on the maturity of the PV cell (Donato et al, 2013). Thus, reduced PV labeling in the cortex might be due to either cell death or impaired maturation of cortical PV cells induced by MPH. Quantification of NeuN (neuronal marker)-positive soma density in different cortical regions in P45 mice failed to reveal reduction in neuronal density (218±8/focal plane in naïve; 255±15/focal plane in MPH, n=6 ctrl and n=5 MPH mice, Mann Whitney test, p>0.5), indicating that we can rule out widespread neuronal loss. To further explore this issue, we employed a transgenic strategy to label interneurons derived from the medial ganglionic eminence, which include PV-positive and somatostatin-positive interneurons (Xu et al, 2008). In particular, we crossed *Nkx2.1Cre* mice with a GFP reporter line. In this crossed mouse line, GFP expression starts at an embryonic stage and is independent of neural activity. We found no difference in GFP+ neuron density in the somatosensory cortex between MPH and control littermates, 5 days following MPH exposure (GFP/NeuN: Layer 23 12±1 in naïve mice and 12.3±0.6 in MPH mice, Layer 56: 25±3 in naïve mice and 21.9±0.9 in MPH mice; 3 mice/experimental group; Mann Whitney test, p>0.5), suggesting that PV cells are still present even if they fail to express the mature marker PV in adult cortex. Of note, we also found no difference in the density of GFP+PV+ cells in P15 MPH vs naïve mice (GFP+PV+/NeuN+: Layer 23 7.3±0.2 in naïve mice and 8.4±0.4 in MPH mice, Layer 56: 9.9±0.9 in naïve mice and 10.3±0.5 in MPH mice; 3 mice/experimental group; Mann Whitney test, p>0.5). These observations suggest that PV cells may slowly fail to increase PV expression during their maturation phase as a result of MPH.

Following our observation on the increase in p75NTR levels following hypoxia, and the parallel decrease in PV expression, we explored the possibility wherein p75NTR may play a specific role in the altered PV cell phenotype found in MPH mice. In order to knockout p75NTR early enough in PV cells, we bred *p75NTR^flx/flx^* mice to *Nkx2.1-Cre* mice and generated both *Nkx2.1-Cre:p75NTR^+/+^* (littermate controls) and *Nkx2.1-Cre:p75NTR^lox/lox^ (cKO)* where p75NTR is ablated embryonically in cortical PV and somatostatin-expressing interneurons. Interestingly we found that *Nkx2.1-Cre:p75NTR^lox/lox^* cKO mice, which experienced MPH at P9, did not show any significant alterations in cortical PV cell density, PV+ intensity (Fig.4B,D; 4G,I; E,J) and memory/spatial recognition (Fig.5C1,D1), contrary to its *Nkx2.1-Cre:p75NTR^+/+^*littermate controls exposed to MPH at the same time. Further, cKO MPH mice showed a better performance in the attentional set shifting task compared to wild-type MPH littermates (Fig.5E1,2), which however did not reach the levels observed in wild-type naïve mice. Overall, these data suggest that p75NTR expression in Nkx2.1Cre derived interneurons makes them vulnerable to MPH, and contributes to the development of specific cognitive dysfunctions.

### Inhibition of p75NTR signaling for one week following MPH rescues long term outcomes on cognition and cortical activity

We next asked whether a pharmacological intervention targeting p75NTR function immediately after MPH would be effective in protecting the neonatal brain from developing long-term cognitive impairments. To this purpose, we administered the small molecule, LM11A-31, which functions as a p75NTR ligand and modulator to downregulate degenerative signaling and to inhibit binding and promotion of degenerative signaling by proNGF (Massa et al, 2006; Shi et al, 2013; Tep et al, 2013). LM11A-31 crosses the blood–brain barrier efficiently when delivered orally or i.p. (Tep et al, 2013), and it has been shown to have protective effects following traumatic brain injury and in Alzheimer’s mouse models (Shi et al, 2013; Tep et al, 2013; Knowles et al, 2013; Yang et al, 2020). LM11A-31, or the vehicle solution, was administered by gavage twice per day for a week following hypoxia exposure (Fig.6A). Adult MPH mice treated with LM11A-31 showed significantly higher memory in the object location and novel object recognition test (Fig.6E1,F1) and performed as well as wild-type mice in the attentional set shifting task (Fig.6G1,2).

**Figure 5.**
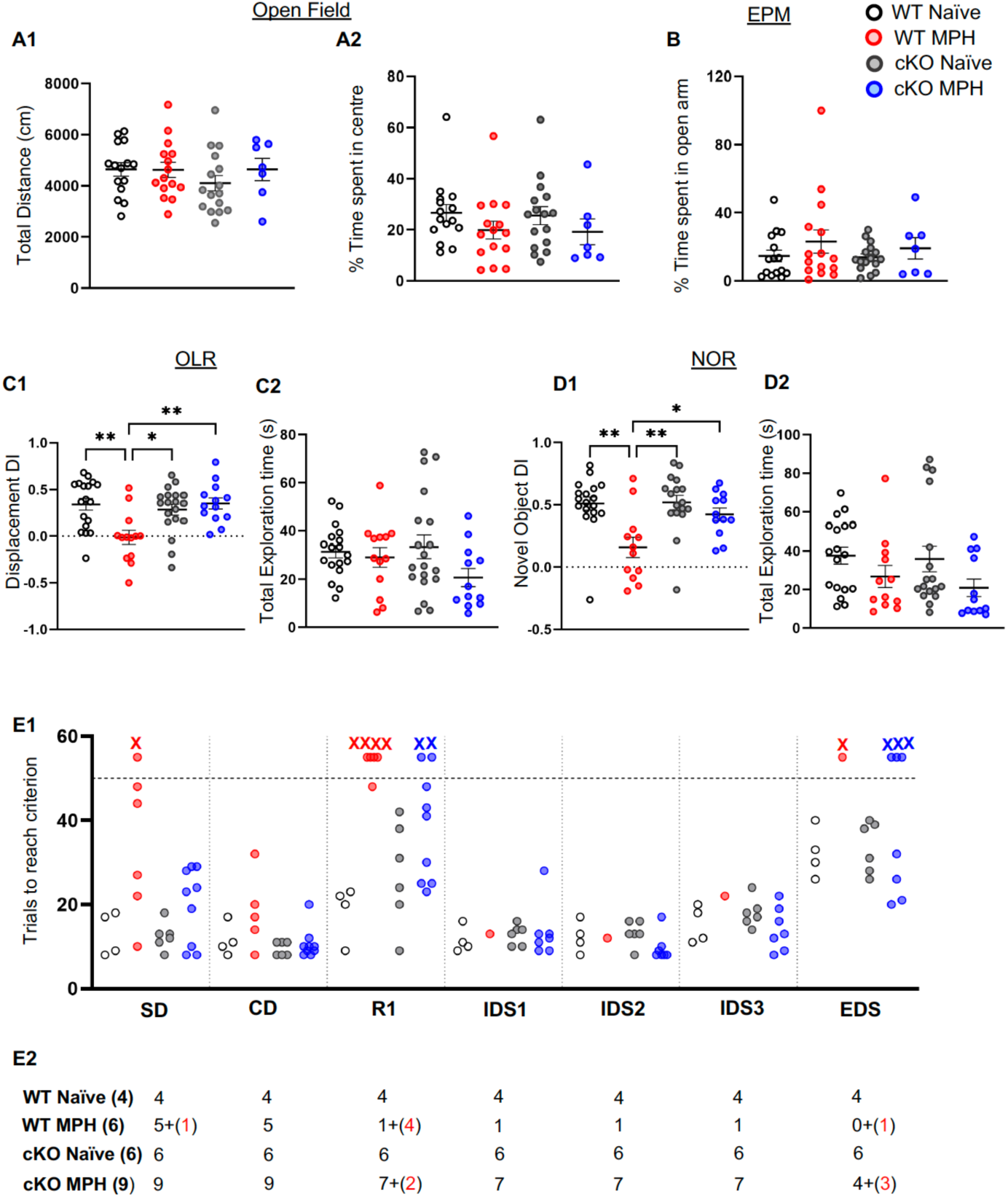
Genetic deletion of p75NTR in MGE-derived GABAergic neurons significantly protects from MPH-induced cognitive dysfunctions. **A1,2.** MPH and naïve mice do not show any significant difference in total distance covered (2-way ANOVA, column p=0.6065; Tukey’s multiple comparisons test; all n.s.) or time spent in the center of the open field (OF), (2-way ANOVA, column p=0.4520; Tukey’s multiple comparisons test; all n.s.). **B.** The time spent in open arms of the elevated plus maze (EPM) is not different between naïve and MPH groups (2-way ANOVA, column p=0.3299; Tukey’s multiple comparisons test; all n.s.). **C1.** cKO MPH mice do not show any memory deficits during object displacement task similar to naïve wild-type or naïve cKO mice, in contrast to wild-type MPH mice while the overall exploration time is similar between all 4 groups (**C2)** (C1, OLR, 2-way ANOVA; Tukey’s *post hoc* multiple comparison test: MPH-WT vs Naive-WT p=0.0085; MPH-WT vs Naive-cKO p=0.04; MPH-WT vs MPH-cKO p=0.0038; MPH-cKO vs Naive-WT p=0.974; MPH-cKO vs Naive-cKO p=0.6848). **D1,2.** cKO MPH mice do not show any deficits during novel object recognition similar to naïve wild-type or cKO mice, in contrast to wild-type MPH mice while the overall exploration time is similar between all 4 groups (D1, NOR, 2-way ANOVA; Tukey’s *post hoc* multiple comparison test: MPH-WT vs Naive-WT p=0.0075; MPH-WT vs Naive-cKO p=0.0032; MPH-WT vs MPH-cKO p=0.033; MPH-cKO vs Naive-WT p=0.9638; MPH-cKO vs Naive-cKO p=0.8643). **E1,2.** MPH mice show a partial rescue in the ASST test, where the crosses represent the mice who did not reach criterion, and failed the ASST task. Red crosses show WT-MPH while blue crosses show cKO-MPH mice. In E2, the number of mice that failed to pass each indicated stage are in red, while those that pass it are in black. Number of mice: **A1-2, B:** Naïve-WT n=15, Naïve-cKO n=16, MPH-WT n=15, MPH-cKO n=7. **C1-2, D1-2:** Naïve-WT n=17; Naïve-cKO n=16; MPH-WT n=12; MPH-cKO n=12. **E1-2:** Naïve-WT n=4; Naïve-cKO n=6; MPH-WT n=6; MPH-cKO n=9. Lines represent Mean ± SEM, while circles represent values for a single mouse.

**Figure 6.**
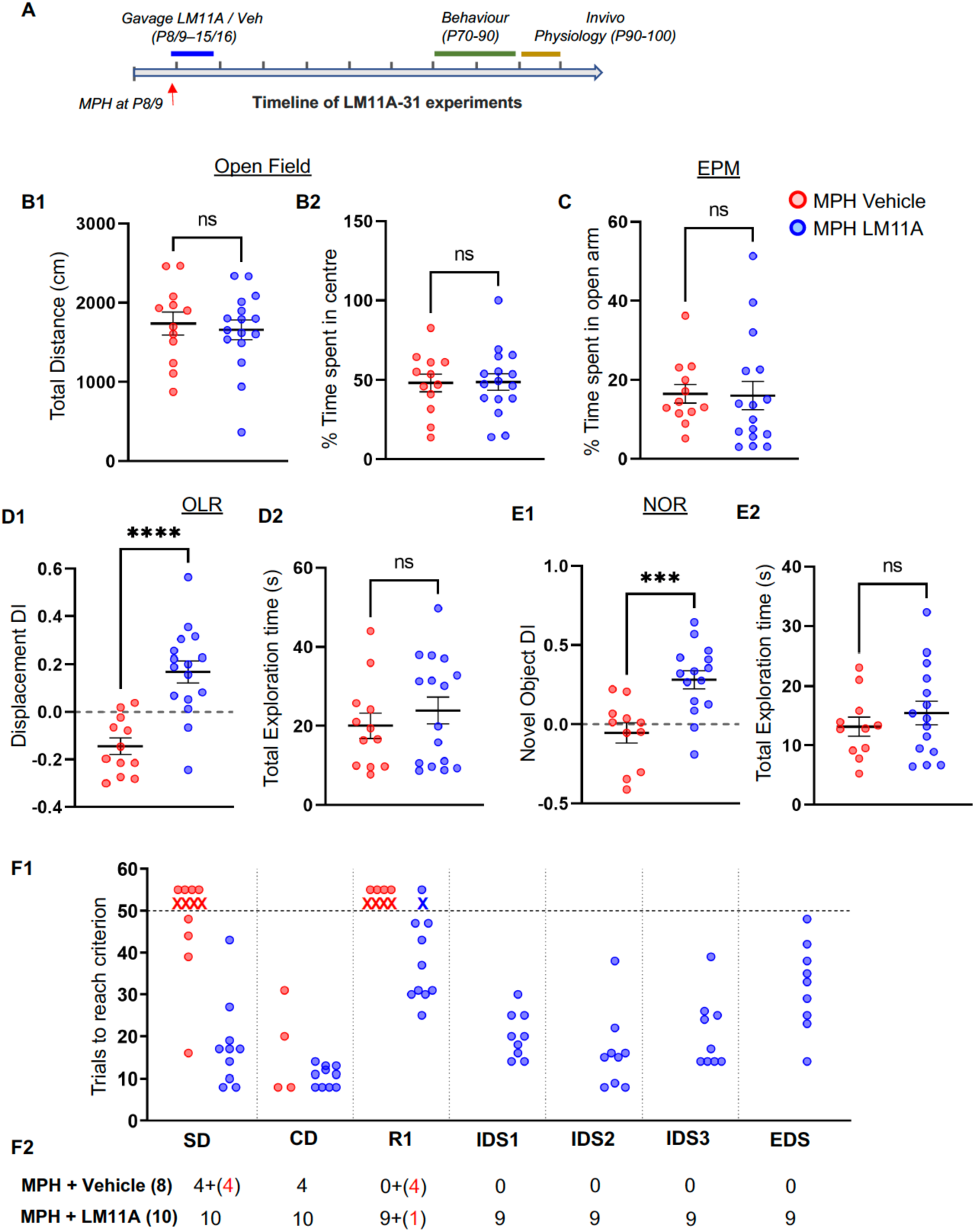
LM11A-31 administration during the first week post MPH prevented the onset of long-term cognitive dysfunctions in adult mice. **A.** Experimental Timeline of LM11A-31 treatment. **B1, 2.** Vehicle– and LM11A-treated MPH mice do not show any significant difference in total distance covered (MW U=88; p=0.73) or time spent in the center of the open field (OF, MW U=95; p=0.98). **C.** Vehicle– and LM11A-treated MPH mice do not show any significant difference in the time spent in the open arms of the elevated plus maze (EPM, MW U=78; p=0.4228). **D, E.** LM11A-treated MPH mice show significant spatial memory (**D1**, OLR, MW U=15; p<0.0001) and novel recognition memory improvement (**E1**, NOR, MW U=20; p=0.0006) as compared to the vehicle treated MPH mice. Number of mice: **B-E.** Vehicle MPH mice n=12; LM11A treated MPH mice n=16. **F1,2.** During ASST behaviour, LM11A-treated MPH mice show a marked improvement in cognitive flexibility. Red cross represents vehicle-treated MPH mice that did not meet the criterion within 50 trials while blue cross presents the same for LM11A-treated MPH mice. In F2, the number of mice that failed to pass each indicated stage are in red, while those that pass it are in black. Number of mice: **F.-G.** ASST; Vehicle-treated MPH mice n=8; LM11A-treated MPH mice n=10. Lines represent Mean ± SEM, while circles represent values for a single mouse.

Neuronal synchronization is a core feature of cortical activity, which is frequently altered in neurological disorders (Uhlhaas and Singer, 2010; Gandal et al, 2012; Fries, 2009). In particular, oscillations in the gamma range are exquisitely regulated by PV cell activity (Fries, 2009; Cardin et al, 2009; Sohal et al, 2009). We thus analyzed gamma power in adult mice during active exploration and found a significant decrease in cortical gamma activity across different cortical regions in MPH vs naïve mice (Fig.7A,B). Conversely, mice treated with LM11A-31 for one week post MPH exposure showed no difference in gamma power compared to naïve mice (Fig.7A,B). Phase amplitude coupling (PAC) analysis further showed reduced coordination of the temporal organisation of oscillatory activity in mice which experienced MPH, which was rescue by LM11A– 31 administration (Supplemental Figure 3).

**Figure 7.**
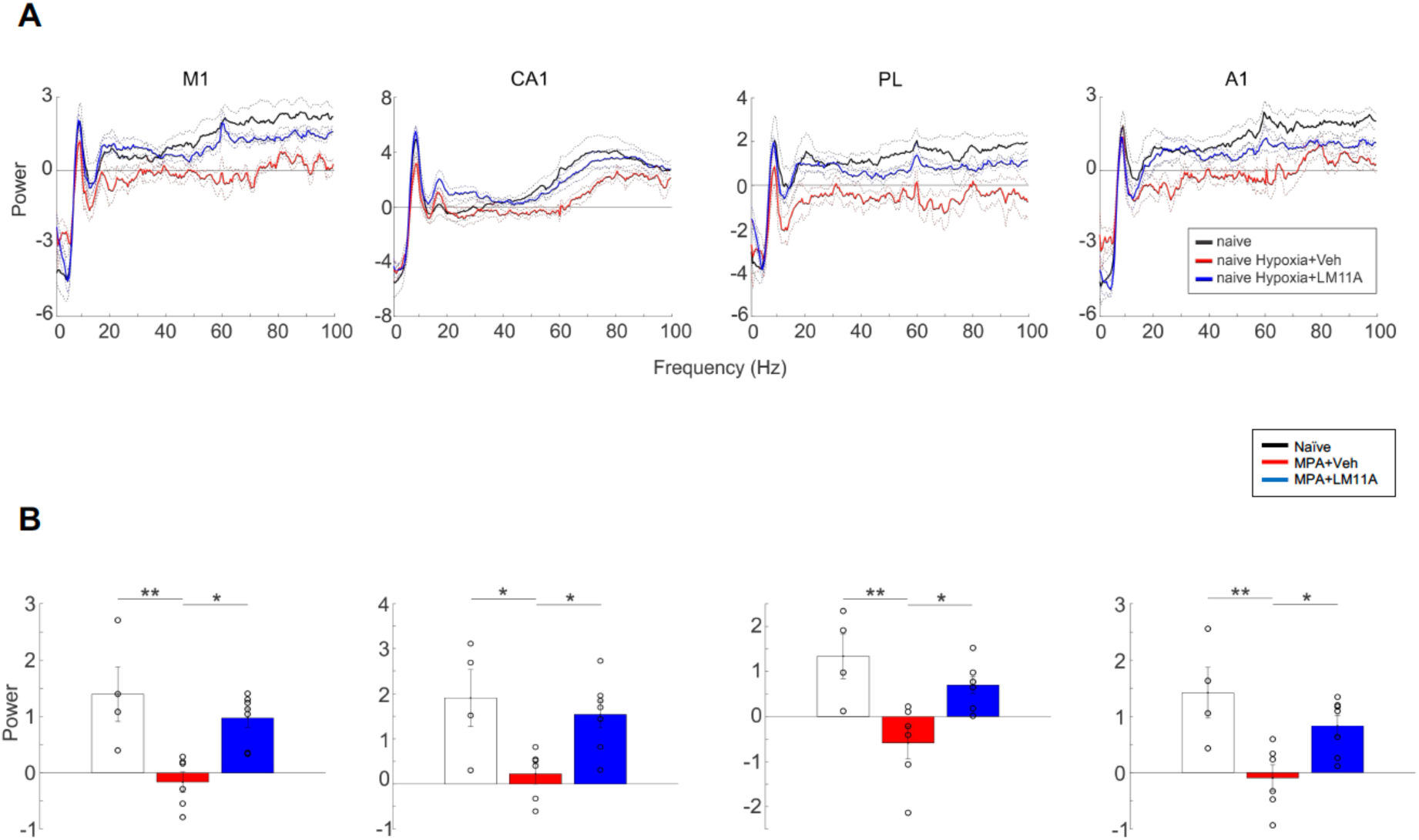
Adult MPH mice show decreased gamma power evoked by active exploration, which is rescued by LM11A-31 administration in the first week post-MPH. **A.** Baseline corrected Power Density Spectra of EEG signal from M1, CA1, PL and A1 during periods of active exploration. B. Bar plots shows mean powers±SEM of 30-80 Hz gamma band in different brain regions. M1: One way ANOVA, F_genotype_ (2,14)=9.6991, p=0.0023, Tukey *posthoc* test showed statistical effect of MPH (naïve vs MPH p=0.0032) and rescue of the drug (MPH + Veh. vs MPH + LM11A p=0.0108 and naïve vs MPH + LM11A p=0.5115). CA1: One way ANOVA, F_genotype_ (2,14)=5.8749, p=0.0140, Tukey *posthoc* test showed statistical effect of MPH (naïve vs MPH p=0.0216) and rescue of the drug (MPH + Veh. vs MPH + LM11A p=0.0365 and naïve vs MPH + LM11A p=0.7779). PL: One way ANOVA, F_genotype_ (2,14)=8.2933, p=0.0042, Tukey *posthoc* test showed statistical effect of hypoxia (naïve vs MPH p=0.0048) and rescue of the drug (MPH + Veh. vs MPH + LM11A p=0.0255 and naïve vs MPH + LM11A p=0.4088). A1: One way ANOVA, F_genotype_ (2,14)=7.4143, p=0.0064, Tukey *posthoc* test showed statistical effect of MPH (naïve vs MPH p=0.0062) and rescue of the drug (MPH + Veh. vs MPH + LM11A p=0.0485 and naïve vs MPH + LM11A p=0.3279). * p<0.05; ** p<0.01). Number of mice: Naïve n=4; MPH+Veh n=6; MPH+LM11A n=7.

In summary, MPH-induced p75NTR upregulation plays a critical role in the development of abnormal brain activity and dysfunctions in specific cognitive domains in adult mice, which experienced perinatal hypoxia.

## Discussion

The main finding of this study is that perinatal mice experiencing a short period (10’) of hypoxia develop abnormal cortical activity, including lower gamma oscillations during active exploration, and cognitive dysfunctions in specific domains as adults. The second main finding is that this perinatal insult causes an increased cortical expression of p75NTR, particularly in PV interneurons, which significantly contributes to the development of long-term neurological and cognitive outcomes.

Correlative, causal and computational evidence indicates that gamma oscillations (30–80 Hz) are strongly modulated by PV cell activity (Fries, 2009; Cardin et al, 2009; Sohal et al, 2009). Since PV expression levels correlate with PV cell activity and maturation state (Donato et al, 2013), the observed decrease in PV intensity in the somata and perisomatic innervations of cortical PV cells is consistent with the findings that gamma oscillations during active exploration are reduced in MPH mice.

p75NTR activation has been shown to activate different pathways, which can lead to apoptosis. However, we did not detect differences in the density of NeuN positive cells in adult MPH mice, which excludes the possibility of widespread cell death. Further, by using a reporter mouse-based strategy, we could not detect any differences in the density of Nkx2.1-expressing cortical GABAergic cells, which include somatostatin-expressing and PV-positive cells, 2 weeks post-MPH. Thus, it is unlikely that increased MPH-induced p75NTR expression leads to increased apoptosis. Instead, we speculate that p75NTR activation may impair the maturation of PV cell innervation by promoting growth cone collapse, via activation of RhoA (Naska et al, 2010) and/or inactivation of Rac signaling, which leads to destabilization of actin filaments and collapse of neurite outgrowth (Deinhardt et al, 2011). Furthermore, p75NTR activation may sensitize neurons to other inhibitory, growth cone collapsing cues, such as Nogo (Yamashita et al, 2003, 2005), ephrins and semaphorins (Lim et al, 2008; Naska et al, 2010). In addition to locally regulating cytoskeletal dynamics, p75NTR activation may cause changes in gene transcription, leading to modulation of expression of PV or/and other proteins regulating PV cell synaptic inputs and/or excitability, which could in turn regulate their maturation state.

Adult naïve *Nkx2.1-Cre:p75NTR^lox/lox^* cKO mice did not show significant differences in the behaviors tested as compared to their naïve control and cKO littermates. In a previous study from our lab, where Cre is expressed under the PV promoter, PV cell-specific p75NTR knockout mice show impaired cognitive flexibility in the same behavioral task we used in this study (Chehrazi et al, 2023). Since there are significant differences in the manner of P75NTR deletion given that a) PV expression starts only after the first postnatal week and peaks just before adolescence in mice, while the Nkx2.1 promoter drives Cre expression embryonically, and b) the Nkx2.1 promoter expresses not only in PV but also in somatostatin-expressing interneurons, it is therefore possible that the developmental age at which p75NTR is deleted differently affects the time course of PV cell maturation, thereby leading to different behavioral outcomes.

Genetic deletion of p75NTR in Nkx2.1-expressing neurons conferred significant protection to the long-term effects of MPH. For example, the density of cortical PV cell somata and the intensity of perisomatic PV puncta in conditional knockout mice that experienced MPH was not significantly different from those observed in naïve wild-type or conditional knockout mice. At the behavioral levels, MPH cKO mice showed normal object and spatial recognition memory. Further, about 50% of MPH cKO mice showed a normal performance in the ASST test, while none of the MPH wild-type mice tested were able to finish this test. Conversely, all adult MPH mice treated with the p75NTR inhibitor LM11A-31 for a week following MPH exposure could successfully complete the ASST test. It is also possible that other cell types are affected by p75NTR upregulation following MPH. In particular, cholinergic neurons located in the basal forebrain express high levels of p75NTR and might be particularly sensitive to hypoxia (Sankorrakul et al, 2021). Impairments in cholinergic transmission could lead to altered attention and cognitive flexibility (Parikh et al, 2020; Prado et al, 2017), and possibly contribute to reduced PV cell activity, which in turn would exacerbate cognitive flexibility and attentional deficits.

In summary, p75NTR upregulation caused by MPH is one critical molecular process contributing to the development of long-term cellular, network and cognitive alterations observed in mice that experience perinatal hypoxia. Future studies should address alternate molecular mechanisms underlying these effects, and explore the contribution of subcortical brain regions.

## Acknowledgements

We thank the Comité Institutionnel de Bonne Pratiques Animales en Recherche (CIBPAR), and all the personnel of the animal facility of the Research Center of CHU Sainte-Justine (Université de Montreal), as well as Dr. Elke Küster-Schöck at the plateforme d’imagerie microscopique (PIM) for instrumental technical support. We thank Dominique Dufour-Bergeron, Josianne Carriço-Nunes, Jean-Francois Toupin, Nathalie Sanon, Pegah Chehrazi and Sebastien Desgent for help with experiments and data analysis. We are grateful to J-F Cloutier for the p75NTR KO mouse tissue and P. Barker for the anti-p75NTR antibody used for western blot.

## Author contributions

BC and GDC designed the experiments. BC, KKYL, MI C-M, AP-R, ML-J and MB performed the experiments. BC, KKYL, MI C-M, AP-R, ML-J and MB analysed data. BC and GDC wrote the manuscript. All authors read and corrected the manuscript.

## Supplemental Figures

**Supplemental Figure 1.**
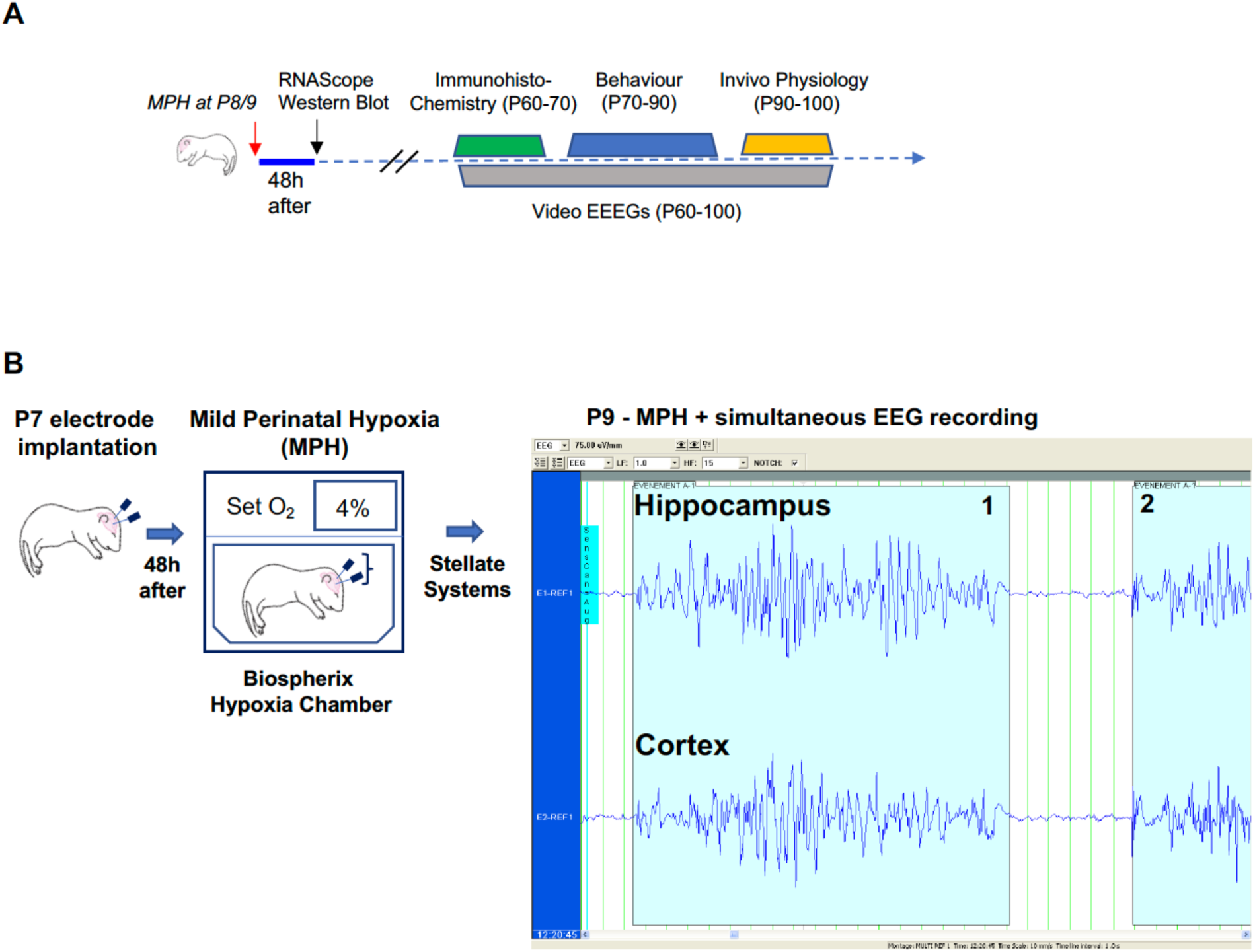
Experimental approach. **A.** Timeline of experimental pipeline following MPH at P8/9. **B.** Schema and example of electrographic hypoxia-induced seizures in P9 mice.

**Supplemental Figure 2.**
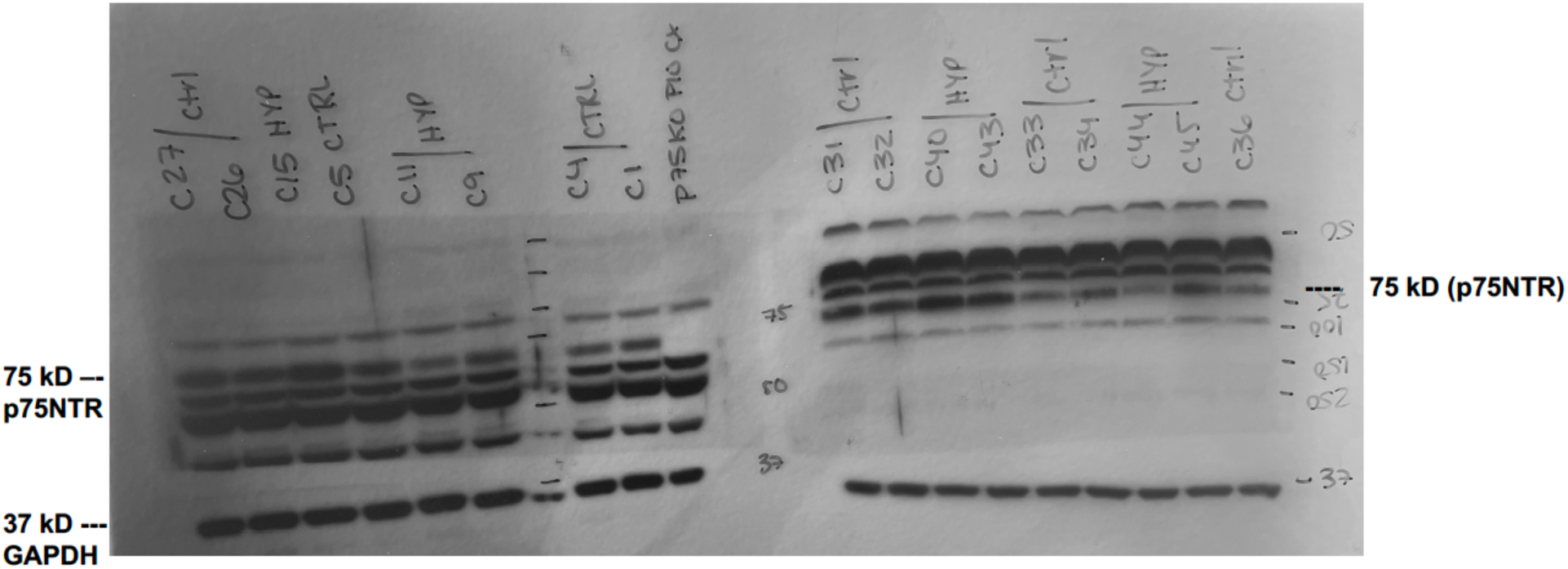
Western blot gels used for data analysis shown in Figure 3. C9 and C11 cortical lysates are from mice with a null p75NTR allele.

**Supplemental Figure 3.**
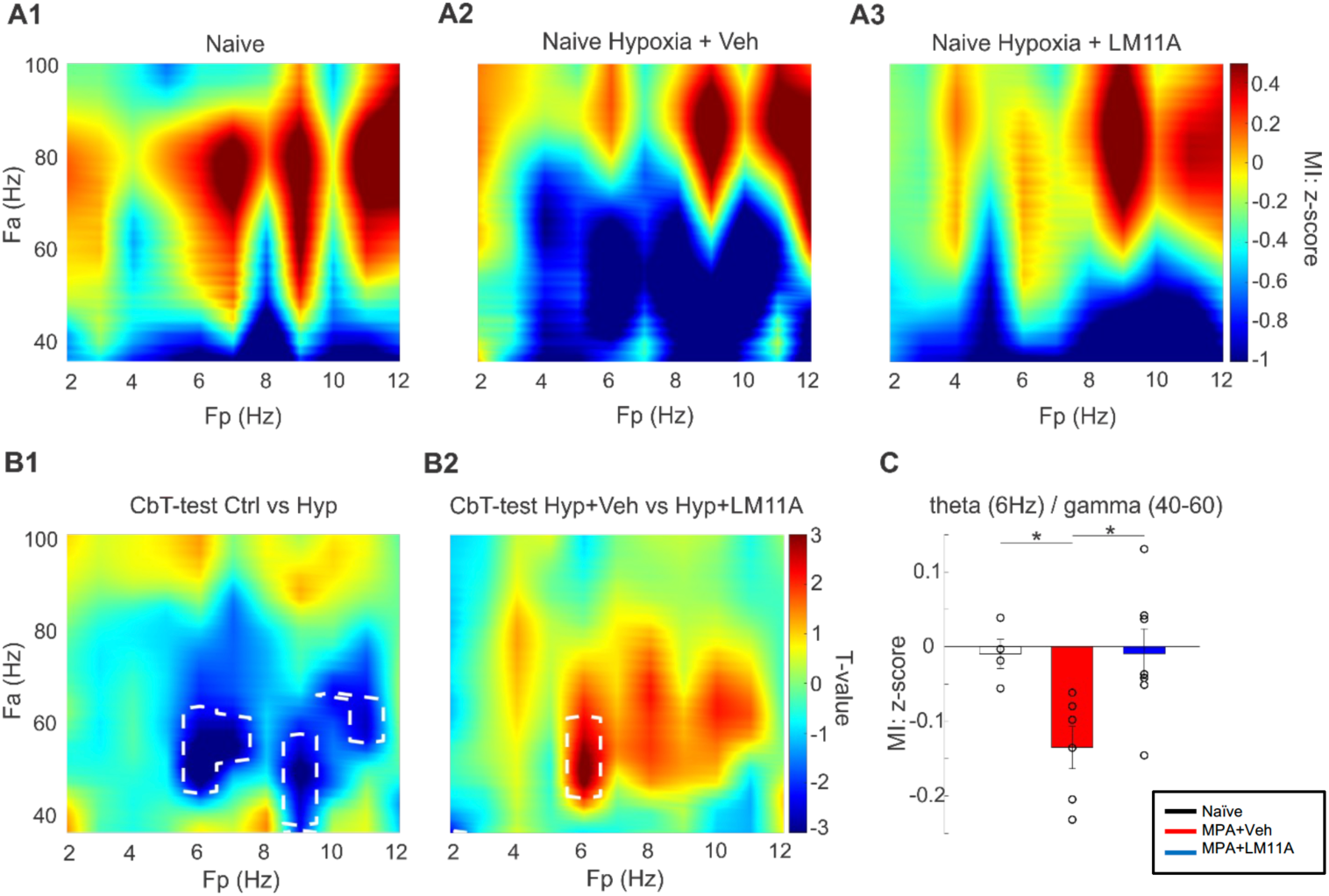
Adult MPH mice show decreased theta/gamma phase-amplitude coupling (PAC) during active exploration, which is rescued by LM11A-31 administration during the first week post-MPH. **A1-A3**. Phase-amplitude co-modulograms in naïve **(A1), MPH** + Vehicle (**A2**) and MPH + LM11A (**A3**). **B1-B2.** Cluster-based statistics comparing PAC modulation index between naïve and MPH + Veh (**B1**) and between MPH + Veh and MPH + LM11A (**B2**). Statistical differences (p<0.05) are marked by white dotted lines. **C.** Bar plot shows z-score values corresponding to theta (6 Hz) /gamma (40-60 Hz) PAC: One way ANOVA, F_genotype_ (2,14)=5.7217, p=0.0153, Tukey *posthoc* test showed statistical effect of MPH (naïve vs MPH, p=0.0469) and rescue of the drug (MPH vs MPH + LM11A p=0.0204 and naïve vs MPH + LM11A p=0.9999). * indicates p<0.05. Number of mice: Naïve n=4; MPH + Veh n=6; MPH + LM11A n=7.

